# Protein Structural Information and Evolutionary Landscape by In Vitro Evolution

**DOI:** 10.1101/582056

**Authors:** Marco Fantini, Simonetta Lisi, Paolo De Los Rios, Antonino Cattaneo, Annalisa Pastore

## Abstract

Protein structure is tightly inter-twined with function according to the laws of evolution. Understanding how structure determines function has been the aim of structural biology for decades. Here, we have wondered instead whether it is possible to exploit the function for which a protein was evolutionary selected to gain information on protein structure and on the landscape explored during the early stages of molecular and natural evolution. To answer to this question, we developed a new methodology, which we named CAMELS (**C**oupling **A**nalysis by **M**olecular **E**volution **L**ibrary **S**equencing), that is able to obtain the in vitro evolution of a protein from an artificial selection based on function. We were able to observe with CAMELS many features of the TEM-1 beta lactamase local fold exclusively by generating and sequencing large libraries of mutational variants. We demonstrated that we can, whenever a functional phenotypic selection of a protein is available, sketch the structural and evolutionary landscape of a protein without utilizing purified proteins, collecting physical measurements or relying on the pool of natural protein variants.

## Introduction

Deleterious mutations can damage the fold and the function of proteins. These mutations are usually rescued, in the course of evolution, by compensatory mutations at spatially close sites that restore contacts and thus preserve structure and function. This creates a correlation between protein contacts and the mutational space of the residues involved that can be compared to shackles. These shackles, that are called evolutionary couplings, can be observed by looking at the covariation between positions in a multiple sequence alignment. Through them, it is possible to predict the network of contacts that determine protein fold. Recently, direct coupling analysis (DCA) and other techniques based on the interpretation of evolutionary couplings have emerged as powerful novel methodologies that enable to predict protein architecture, fold and interactions (Weigt et al. 2009; Marks et al. 2011; Morcos et al. 2011; Ekeberg et al. 2013; Kamisetty et al. 2013; Ovchinnikov et al. 2014; Ovchinnikov et al. 2017). These techniques have immensely increased the arsenal of tools at the scientists’ disposal to obtain structural information (Altschuh et al. 1987; Göbel et al. 1994; Pazos et al. 1997). One of the several advantages of an evolution-based approach is also the possibility to obtain structural information of proteins notably difficult to crystalize and/or model, such as membrane (Hopf et al. 2012) or disordered proteins (Toth-Petroczy et al. 2016). DCA has been successfully applied at the proteome scale leading, for instance, to the successful prediction of all the binary protein interactions in *E. coli* (Hopf et al. 2014) and the retrieval of the structures of entire protein families and subfamilies present in the PFAM database (Uguzzoni et al. 2017).

We wondered if these tools could be applied to other types of evolutionary data such as libraries of proteins evolved *in vitro* by carefully controlled mutations and selection (Chen and Arnold 1993; Zaccolo and Gherardi 1999). This artificial form of evolution generates a collection of functional variants of the protein of interest by coupling a targeted mutagenesis of the gene to a strong selection pressure for the desired phenotypic trait. The method is widely used in synthetic biology as a tool of protein engineering. It is also important in studies aimed at understanding evolutionary pathways. The application of DCA on an artificial library would give the possibility to generate data without the need of relying on natural evolution, paving the way for structure determination by artificial selection in vitro. This process is however very challenging, because the construction of molecular evolution libraries requires a platform able to sequence the whole gene, at the risk of losing the co-occurrence of mutations in distant positions. Another constraint lies in the size of the mutational space sampled by molecular evolution because coupling techniques need a high-complexity highly-mutated collection of sequences to retrieve couplings.

Here, we describe a general methodology based on molecular biology techniques coupled to computational analysis. Our method goes all the way from an original ancestor gene sequence, to the generation and collection of sequences, to data analysis using molecular evolution. We generated a large library of variants of a target gene, followed by in vivo phenotypic selection to isolate functional variants of the ancestor protein. The plasmid library carrying the mutants was then sequenced and analyzed by DCA. By this method, we were able to demonstrate that we can retrieve evolutionary constraints and get partial information on protein structure. During the course of this artificial evolution of an ancestor gene, we noticed that the sequences collected after cumulative rounds of mutagenesis become progressively more similar to the collection of natural variants. They are ultimately comparable to an early stage of the natural evolution of the protein, when the variants explored are still fairly similar to the founding progenitor that underwent mutagenesis.

By substituting natural with *in vitro* evolution, we explored a brand new application of DCA which overcomes the limitations that have so far hindered the generality and scalability of the method. As a proof of concept, we chose TEM-1 beta-lactamase, a member of the Beta lactamase family of enzymes that confer to bacteria the ability to destroy the beta lactam ring of penicillinand derivatives such as ampicillin (Abraham and Chain 1940). Resistance allows bacteria to grow in the presence of these antibiotics, a function that is easily amenable to a phenotypic selective pressure. TEM-1 is a golden standard for molecular evolution experiments (Bershtein et al. 2006; Salverda et al. 2010; Deng et al. 2012; Jacquier et al. 2013; Firnberg et al. 2014; Stiffler et al. 2015). Our data clearly demonstrate that proteins evolved by molecular evolution can be used to collect evolutionary and structural data and provide a new tool to all branching fields of evolutionary coupling and molecular evolution research.

## Results

### Experimental design

We employed random mutagenesis from error prone PCR (Wilson and Keefe 2001) to generate a large library of variants of the target gene, followed by transformation into bacterial cells and *in vivo* phenotypic selection to isolate functional variants of the ancestor protein (**Figure 1A**). The plasmid library carrying the mutants was then collected from the surviving bacteria and subjected to Pacific Bioscience single molecule real time (SMRT) sequencing (Eid et al. 2009). We used the TEM-1 beta-lactamase of the pUC19 plasmid (Norrander et al. 1983). TEM lactamases are encoded by genes around ~900 bp in length and are present in several natural variants (Bush 1997). Their structure consists of a three-layer (αβα) sandwich (**Figure 1B**). As a reference for the mutational landscape, a collection of beta lactamase sequences (named “UniProt” dataset) was obtained from the UniProt database. To obtain a heavily mutagenized beta lactamase without damaging the survival rate, the library was subjected to consecutive cycles of mutations, selection and amplification through the use of error prone PCR (Wilson and Keefe 2001) and growth in selective semisolid media (Elsaesser and Paysan 2004; Fantini, Pandolfini, et al. 2017).

**Figure 1:**
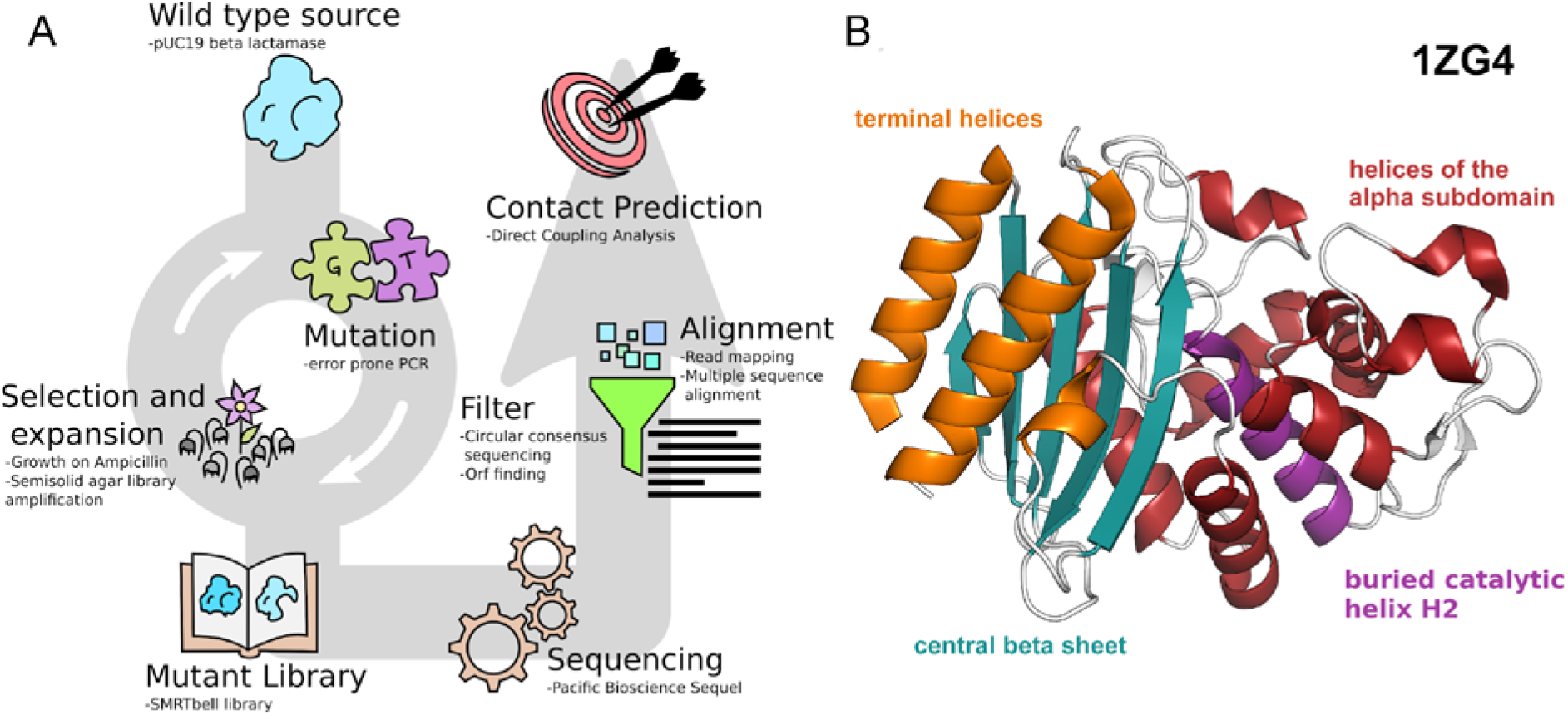
Schematic representation of the pipeline and structure of the target protein. A) The coding sequence of target protein is cloned in a plasmid vector for mutagenesis. After several rounds of mutation and selection for the desired function, the new protein variants are collected in a DNA library. NGS of this library provides sequences that after processing are used for generating the prediction. B) Experimental structure and main features of TEM-1 beta lactamase (derived from PDB 1ZG4). The N- and C-terminal helices (orange) form a subdomain with the five stranded central beta sheet (blue) linked to the helical subdomain by two small hinge regions (red) on the opposite side of the sheet. The catalytic pocket resides at the interface between the beta sheet and the helical domain. Helix H2 (purple) is the innermost helix of the helical domain and comprises both a catalytic and a structural function.

Classical directed evolution performs the selection process in solid media and the results is usually limited to few tens of thousands colonies directly proportional to the number of Petri dishes employed. Cultures grown on solid media are not easily scalable and the biomass they produce is limited. On the other hand, liquid media cultures are easily scalable and produce a large amount of biomass but fail to preserve the library complexity and distribution. In liquid media, fast growing phenotypes are not constrained and thus tend to dominate the culture while rare variants are prone to disappear. Maintaining a high complexity is critical, so neither solid nor liquid culture are the optimal solution. The issue was bypassed by encapsulating the colony forming units (CFUs) able to survive the selection in a matrix of a semisolid medium which allows local growth but prevents diffusion (**Suppl Figure S1**). After colonial growth the plasmid library can be collected from the media by centrifugation. We will refer hereafter to the library at the end of each cycle as a generation of molecular evolution. In total, we performed twelve generations. The first, fifth and twelfth generations were sequenced with the Pacific Bioscience (PacBio) Sequel platform and analyzed. Deep sequencing is able to sequence millions of reads.

### Molecular evolution libraries mimic natural variability

We first thought to perform complete combinatorial two-residue deep mutational scanning to create a library (Olson et al. 2014). However, although powerful, deep mutational scanning does not represent the mutational space that nature would explore during evolution. We used instead error prone PCR to drive mutagenesis and a phenotypic selection to collect the functional variants to mimic natural evolution. To survive the selective environment, the bacterial cells had to incorporate one of the plasmids of the mutant library. The variant of beta lactamase carried by the plasmid had to maintain functionality after mutagenesis. The first event is favored by a high transformation efficiency allowed by the use of the pUC19 plasmid vector, while the enzyme functionality is expected to decrease with the incremental number of mutations introduced in the lactamase sequence. The mutations generated during mutagenesis are at the same time necessary for evolution but harmful for survival. To obtain a high mutational load while still guaranteeing a good amount of survivors we applied a generational approach, where new mutations were built on a collection of mutated but functional sequences from the previously selected generations. The number of mutations and the related final 2.5 – 3% survival rate was regulated by limiting the error prone PCR to 20 cycles every generation. To verify the progress of molecular evolution and maintain libraries with a fair amount of complexity, we controlled three parameters throughout twelve generations: the number of transformants in the bacterial growth, the number of mismatching amino acids in a small sample of clones and the information entropy at each amino acid position.

Since each bacterial colony in the selection medium expresses a single functional variant of the protein, the number of transformants poses a theoretical upper limit to the library diversity. We kept the number of transformants at least in the same order of magnitude of the sequencing capacity of the next generation sequencing (NGS) platform (between 100,000 and 1 million) to guarantee a good library complexity (**Suppl Figure S2**). We raised this limit to 400 thousand clones in the last few generations to increase the probability to sequence unique variants. After each generation a small sample of clones underwent sequencing to estimate the number of mismatching nucleobases and amino acids with respect to the ancestor sequence (**Figure 2A**). After 12 generations of molecular evolution when sequences had a median of 25 mutations in the peptide chain, the system is still showing a nearly-linear increment in the number of mutations per generation.

**Figure 2:**
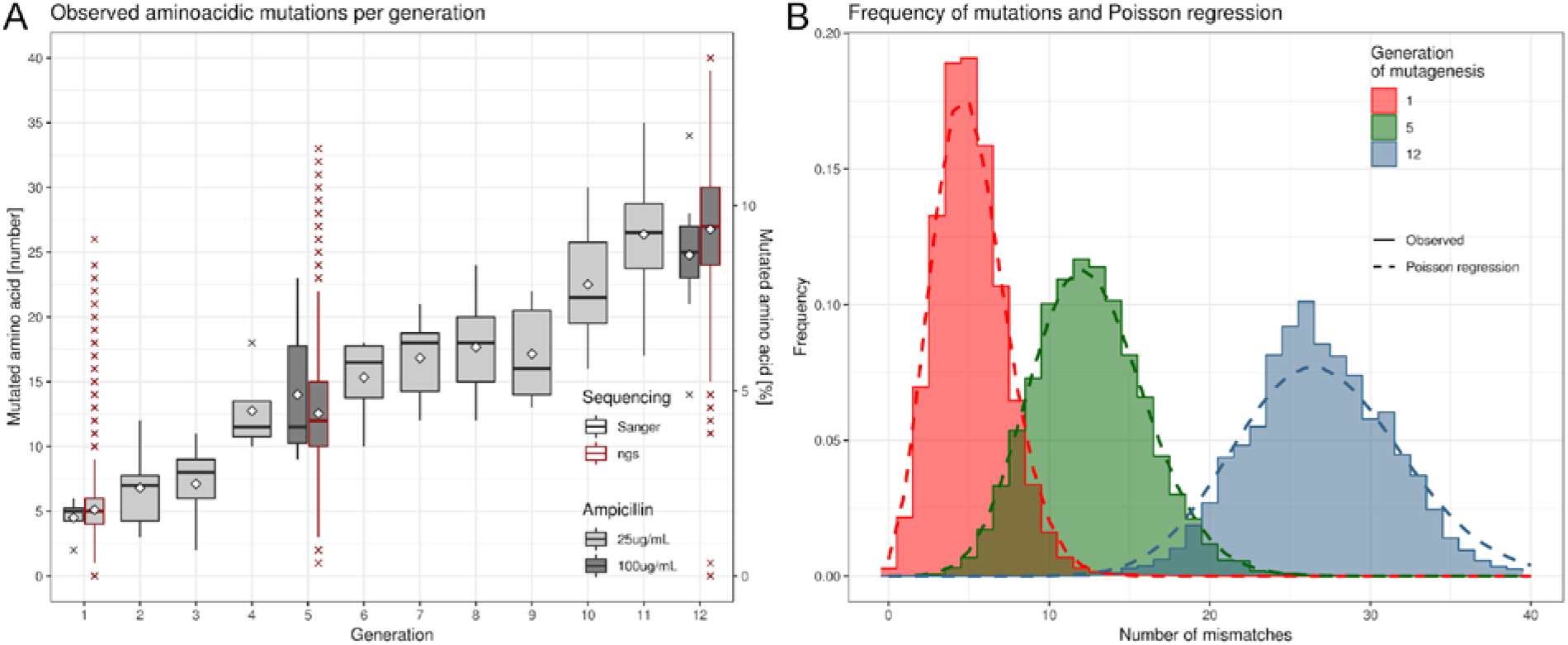
Sequencing and molecular evolution results. A) Boxplot showing the number of amino acid mutations (mismatches) observed in the sample of clones sequenced after each generation (Sanger sequencing, black border) and after NGS (red border). The white diamond dots indicate the mean. B) Frequency distribution of the number of aminoacidic mutations observed in the sequenced libraries (solid lines) and their respective Poissonian regressions (dotted lines: gen1: λ=5.12; gen5: λ=12.54; gen12: λ=26.9).

To complement this information, the same parameter was estimated from the sequencing results of the three sequenced generations. The distribution of the number of mismatches per sequence fitted the theoretical Poissonian model expected for a mutagenesis (gen1: λ=5.12 s=0.0054; gen5: λ=12.54 s=0.0084; gen12: λ=26.9 s=0.0159) (**Figure 2B**). The median number of mutated residues observed when the colonies were picked matched perfectly that obtained from NGS (**Figure 2A**) and what was expected from a Poissonian model, proving that the handful of colonies picked are representative of the mutations present in the library. We concluded from the observed steady increase in the number of mutations throughout molecular evolution that the final mutation fraction of the evolved protein library can be regulated by increasing the number of generations. Sequencing data also allowed us to calculate the mutation fraction per amino acidic position, defined as the frequency of the observed mismatching amino acids compared to the original pUC19 beta lactamase sequence. After 12 generations of molecular evolution we started to observe several instances of genetic drifts, in which a mutation became more common than the original residue at a given position (**Suppl Figure S3**). This phenomenon makes the mutation fractions less informative, since they involve a comparison to the original residue that is now a minority. To circumvent the problem, we measured the Shannon information entropy of each residue, obtaining an approximation of the mutagenesis impact for each position, without the need of a reference sequence (**Figure 3A, Suppl Figure S4**). The proportion of mutants and the information entropy of each residue were strongly correlated to each other and with those observed from the UniProt dataset (mutant frequency: rho 0.624, p < 1e-15; entropy: rho 0.632, p < 1e-15) (**Suppl Figure S5**). We also observed that, at each position, both the entropy and the mutation frequency of the molecular evolution libraries are almost always lower than the corresponding ones from natural evolution (**Figure 3B, Suppl Figure S5**). This is likely a limit to which a molecular evolution library would tend, given enough mutagenesis rounds.

**Figure 3:**
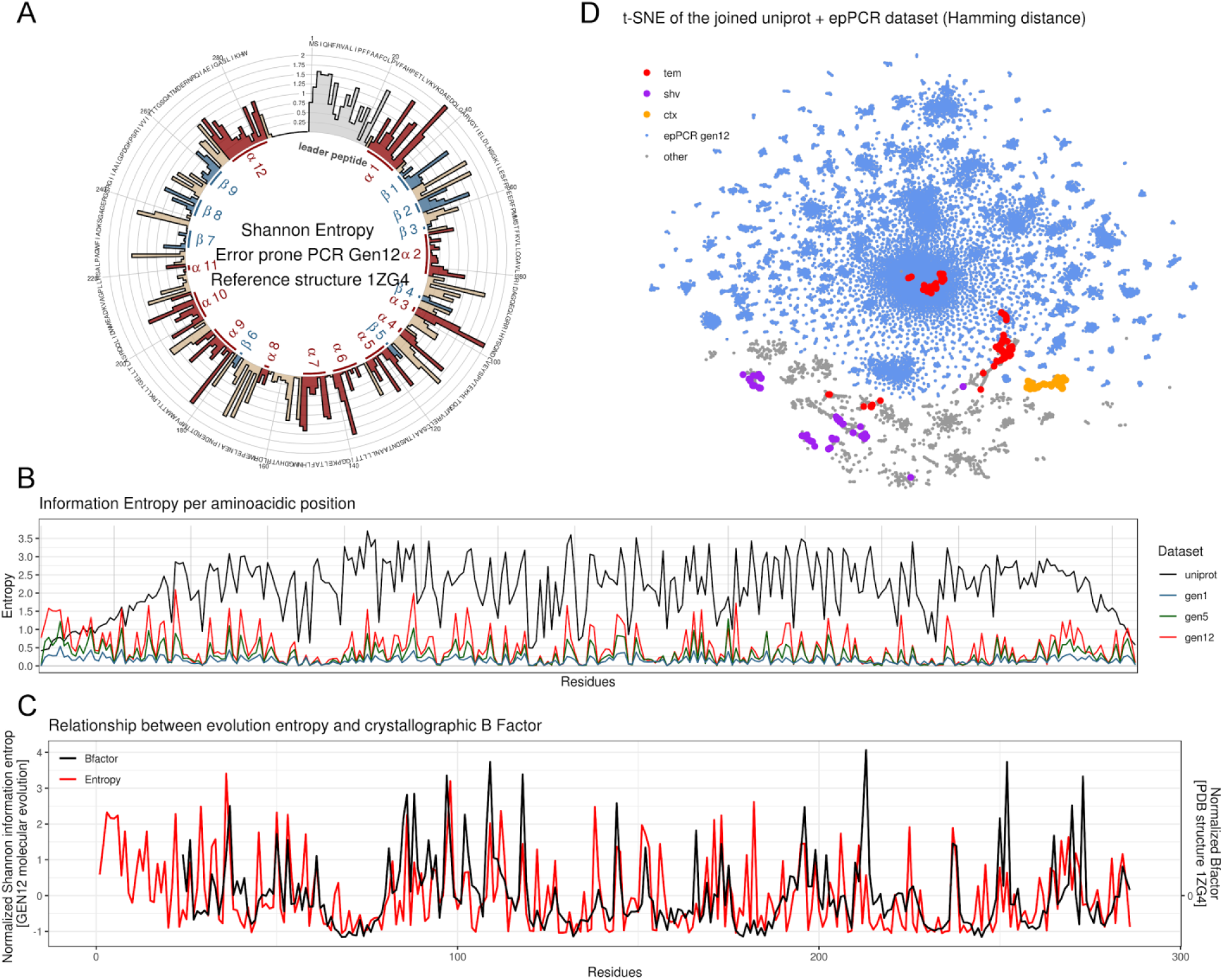
Sequencing and molecular evolution results. A) Shannon information entropy (H) per residue position of the sequenced 12th generation library. The colors and annotations follow the secondary structure classification present in the PDB structure 1ZG4 (red: alpha helices, blue: beta sheets, tan: coils). The leader peptide sequence (light gray) is missing in the structure. B) Comparison of the Shannon information entropy between the UniProt and the in vitro evolved datasets. C) Relationship between the entropy of the residues obtained in molecular evolution and the mean B factor of the residues observed in the reference structure 1ZG4. Since the reference structure is missing the leader peptide, the first 23 amino acids do not have an associated Bfactor. D) t-SNE dimensionality reduction applied to the joined UniProt / error prone PCR 12th generation library dataset. Hamming distance between sequences was used as distance metric. Gray and cyan represent the original dataset (gray UniProt, cyan epPCR library). Overlaid on top, the UniProt sequence membership to one of the three main families of type A beta lactamases retrieved from the corresponding UniProt annotation are displayed in bright colors. The original pUC19 beta lactamase before molecular evolution is classified as a TEM beta lactamase (red).

For reference, the Ostermeier’s mutational database from deep mutational scanning experiments on TEM-1 has a similar entropy profile (Firnberg et al. 2014). We constructed a list of viable mutations by pooling together the mutations from Ostermeier’s data able to grow in 32 μg/mL ampicillin and above in a single collection. The Shannon entropy of the Ostermeier’s database correlates with all our sequenced generations and in particular with the first generation (correlation with gen1: rho 0.560; gen5: rho 0.530; gen12: rho 0.494; all of these with a p value < 1e-15). In the first generation, the accumulation of mutations is reduced and the network of interconnected residues is less developed compared to the last generation. Thus our entropy profile is more similar to what can be obtained from a single residue deep mutational scanning.

### Single molecule sequencing overcomes library restrictions

We used the PacBio single molecule real time (SMRT) sequencing platform (Sequel) (Eid et al. 2009) that can obtain up to a million readings per sequencing cell (van Dijk et al. 2018) and is compatible with the complexity of a molecular evolution library. The total number of transformants for the three sequenced libraries, that pose a limit to the library complexity, were 200K, 260K and 400K CFUs, respectively, while each sequencing run generated 192K, 289K and 157K raw readings after quality filtering. The sequenced DNA fragment was over 800 base pairs. Other more common NGS platform like Illumina HiSeq or MiSeq are instead characterized by decreasing quality with increasing base position (Kircher et al. 2009) and thus cannot sequence more than few hundreds base pairs. It is possible to scale up the number of sequencing cells, the amount of bacteria that undergo the selection process and the size of the DNA fragment used in mutagenesis to satisfy the requirement of this technique for any desired protein. Our mutational library is the first molecular evolution library sequenced in a third generation sequencer, thus guaranteeing a high volume of high quality single molecule data. This library is also one of the most mutated TEM beta lactamase libraries ever produced, where its elements diverge from the ancestral protein for ca. 10% of their original amino acidic composition.

### The mutational landscape of the evolved library reflects the structural features of TEM beta lactamases

The beta lactamase structure (PDB entry 1ZG4, Stec et al. 2005) was used as a reference structure to assess the contact predictions and the accuracy of the analysis. TEM1 beta lactamase is a globular protein with a roughly ellipsoidal shape (**Figure 1B**) (Jelsch et al. 1993). It can be divided into two subdomains. One is composed of a five stranded beta sheet, the N-terminus and the two last C-terminal helices. The second is a big helical subdomain located on the other side of the sheet. The protein contains a large hydrophobic core between the beta sheets and the helical subdomain, and a second hydrophobic region in the core of the helical domain. The innermost helix of this domain, H2, contains both structural and catalytic residues. The PDB structure lacks the first 23 amino acids, corresponding to the leader sequence for secretion, which is cleaved during protein maturation to allow protein release.

The profiles of the mutation rate and entropy per residue observed in our molecular evolution libraries are conserved and increase across generations, in line with what is observed in the UniProt dataset of the naturally evolved beta lactamase family (**Figure 3B**). This profile reflects the different mutation propensities of the various residues as well as the interactions with the solvent and the polarity of the local environment. We observed a high degree of conservation in the presence of bulky nonpolar amphipathic residues like tryptophans (W2108, W227 and W286) and methionines (M184, M209, M268) and in cysteines involved in the sulfur bridge (C75, C121), whilst small residues show in general an increased variability (**Suppl Figure S6**). Small nonpolar amino acids such as valine, leucine and isoleucine form a group of interchangeable residues in several positions (45, 54, 171, 196 and 244). The small polar counterparts glutamate and aspartate can be found replacing one another in others (33, 36, 113, 195, 269).

A periodic alternating pattern of high and low entropy can be seen in the long alpha helices H1, H9, H10 and H12. This reflects the nature of the two halves of the helices, one being hydrophilic partially exposed to the solvent, the other containing hydrophobic residues packed against the protein core. H2 is different from the other helices because it is located deeply inside the hydrophobic core of the protein and mediates most of the hydrophobic interactions of the protein. This parallels the lower mutation frequency and entropy observed in all our libraries (**Figure 3A**, **Suppl Figures S3 and S4**), since mutations in the hydrophobic core have a high chance to damage the fold and thus impair protein function.

It is also noteworthy the correlation (Spearman correlation: rho 0.53, p < 1e-15) between the mean crystallographic B factors of residues in the reference structure and the information entropy retrieved from the evolved library (**Figure 3C**). This correlation likely reflects the tendency of residues that are part of ordered structures to be averse to mutation.

While the mutational landscape of TEM-1 beta lactamase covers a broad range of substitutions, four mutations became more frequent than the original sequence in the last generation of molecular evolution by genetic drift: M180T, E195D, L196I, S281T (**Suppl Figure S3**). Among these, M180T (M182T in the standard numbering scheme of class A beta lactamases, Ambler et al. 1991) is a well-documented mutation known to contribute to the protein stability and found both in natural variants (Huang and Palzkill 1997; Wang et al. 2002) and in mutagenesis experiments (Goldsmith and Tawfik 2009). E195D and L196I (E197D and L198I in standard numbering) are mutations in the H8/H9 turn which are commonly found during mutagenesis (De Visser et al. 2010). D197 is the consensus amino acid for this position (197) in the original alignment of class A beta lactamase (Ambler et al. 1991).

We next used principal component analysis (PCA) on the Shannon entropies associated to each position of each dataset, to evaluate the evolution of the mutagenized libraries towards the natural diversity (**Suppl Figure S7**). We also applied PCA (**Suppl Figure S8**) (Wang and Kennedy 2014) and t-SNE (**Figure 3D**) on the sequences themselves, to evaluate the degree of dispersion for each generation in comparison to the natural variants. These analyses suggest that subsequent mutagenesis cycles consistently evolve the sequences in a concerted direction that is similar to that observed in the natural dataset. t-SNE also suggests that the cluster of the evolved lactamase is only an extension of the TEM family and does not cluster with other members of class A beta lactamase (**Figure 3D**). Thus the molecular evolution libraries describe the mutational space of a specific protein and not of a protein family.

From this analysis we may conclude that the library has retained the most salient characteristics of natural beta lactamase variants and exclusively represents the mutational landscape around the protein of interest. This means that the library provides a snapshot of the early stages of evolution, neither too similar nor too diverse from the original sequence, but exploring the landscape of mutational substitutions in a direction analogous to that followed by natural selection.

### The observed mutational events mimic the early stages of protein folding

After several generations, we extracted the longest open reading frame from each of the 157K circular consensus reads obtained after sequencing the last generation of mutagenesis and removed those shorter than the wild type protein. We built a multiple sequence alignment (MSA) from the remaining 106K (68.9%) translated peptides and kept only the original 286 positions related to the wild type enzyme. To predict which residue pairs interact, we applied a custom implementation of DCA that applies a pseudo-likelihood approximation (Balakrishnan et al. 2011) to this MSA as well as to the MSA obtained similarly from the other two sequenced generations of mutagenesis (see Materials and Methods). We retained the 286 residue pairs (0.72% of the total possible contacts) which showed the highest DCA score and were more than five residue apart in the MSA and compared them to the contact map of the reference structure (gen1: **Suppl Figure S9**, gen5: **Figure 4A**, gen12 **Figure 4B**).

**Figure 4:**
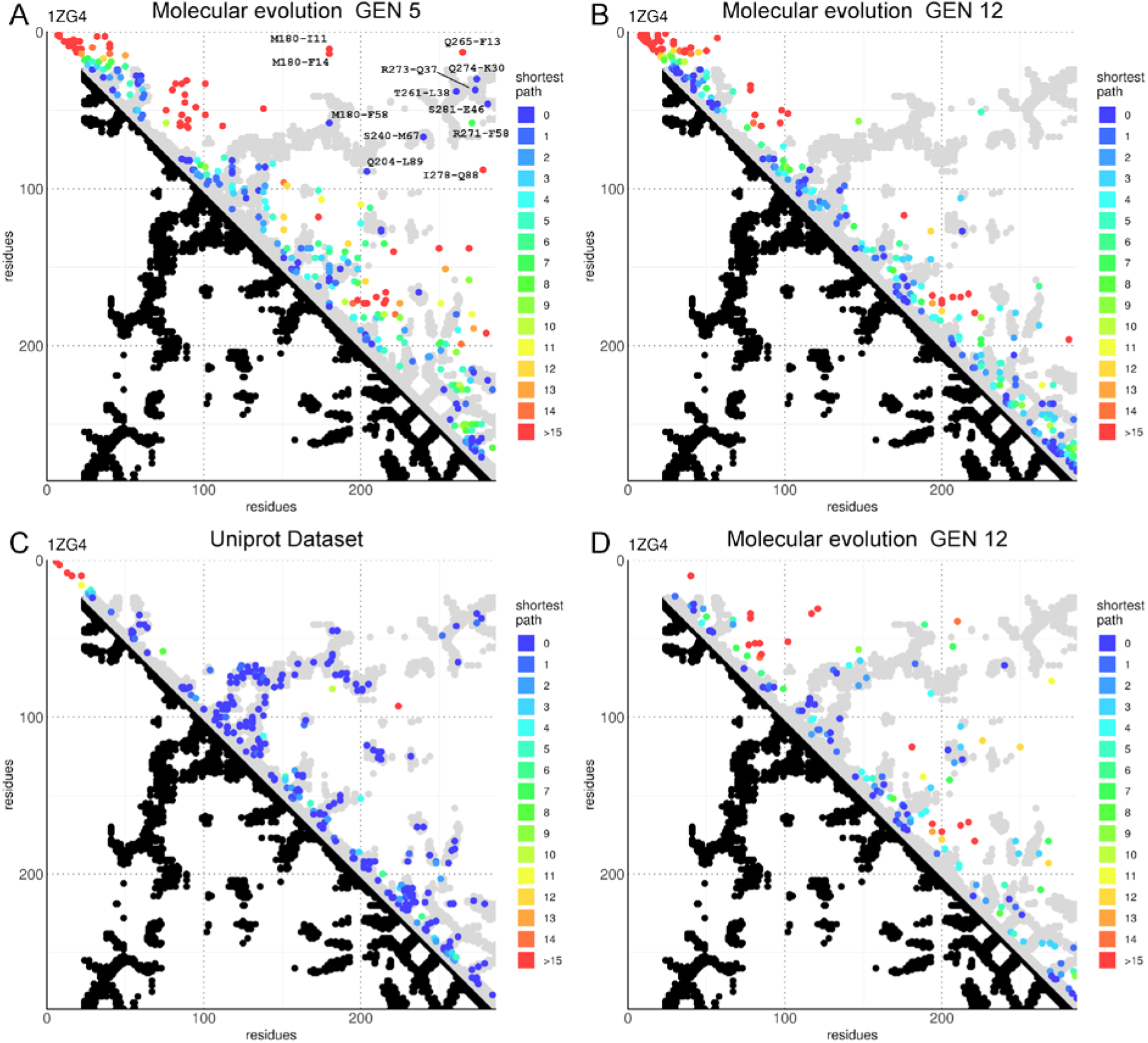
DCA predictions of the beta lactamase contact map. A) DCA plot showing the top L (L = 286, the length of the protein amino acid chain) contact predictions by DCA obtained from the fifth generation of molecular evolution. The graph is an LxL grid where each axis represents the amino acid positions of the lactamase chain, from the N-to C-terminals. Each point represents the pair of residues described by its coordinates. The graph is separated in two halves. In the lower half black dots represent pairs of residues that have at least a pair of their respective non-hydrogen atoms less than 8.5 Å apart in the reference crystallographic structure (PDB id: 1ZG4). These positions are considered residues in contact with each other. In the upper half the top L DCA predictions from the molecular evolution dataset are plotted above the gray mirrored silhouette of the crystallographic contacts. Pairs where the respective residues are less than 5 positions apart in the lactamase alignment are excluded from this ranking to promote visualization of long range interactions. In the graph the color indicates the shortest path (as the lowest L1 norm in the graph grid space) connecting the point to a contact pair position (a pair of residues that have non-hydrogen atoms less than 8.5 Å apart in the reference structure). B) DCA plot showing the top L (L = 286, the length of the protein amino acid chain) contact predictions by DCA obtained from the 12th generation of molecular evolution. C) Plot of the top L DCA predictions of the UniProt dataset. D) Plot of the top L/2 partial correlations of residue positions on DCA score obtained from the 12th generation of molecular evolution.

The first generation library was clearly unable to provide any meaningful result, while the fifth generation showed an interesting pattern. We may observe that the predictions of interacting residues made from the fifth generation library tend to cluster and are crowded in the area near the diagonal. Of particular interest are the elements at the N-terminus (residues 1-60) where the prediction clearly overlaps with the interactions made by the first few N-terminal secondary structure elements (helix H1 with the first two strands of the central beta sheet). Other important clusters can be seen in correspondence to the branching points from the diagonal (near the diagonal around residues 100, 160, 200 and 260). This set of contacts running perpendicular to the diagonal reflect the presence of hairpins. The clustering at the branching point should be expected because the branching point is where the chain inversion takes place. Since loops are more flexible and solvent exposed than other secondary structure elements (Schlessinger and Rost 2005), their composition is less critical for protein fold and allows a broader range of variation. More variations in amino acid composition increase the probability to observe a covariation pattern in DCA. Long range contacts, the most important interactions to reconstruct the tertiary structure of a protein, were also observed (**Figure 4A**). Most of them overlap with the structural trace, with only five exceptions that do not match the reference (M180 is the position that shows the strongest genetic drift and is very noisy). Two of these long range contacts predict the interactions between the N- and C-terminal helices (Q274-K30, R73-Q37). The other two mediate the interaction of the terminal helices with the beta sheet: T261-L38 mediate the interaction between the middle strand B9 of the sheet and the N-terminal helix H1; S281-E46 mediates the interaction between the C-terminal helix H12 with the first beta strand B1. M180-F58 is a contact between the loop between B6 and H9 and the lateral beta strand B2. This is an interesting area because F58 stands at the very beginning of one of the two hinge regions that connect the two domains of the protein. The last two contacts are at the opposite ends of the innermost helix H2 that carries both a catalytic and structural function. Q240-L89 is an interaction between H10, the last helix of the helical domain, and the loop at the end of H2, while S240-M67 mediates the interaction of H2 with a conserved serine in the loop between the beta strands B7 and B8.

The data are still too sparse to define clear cut interaction zones and tend to cluster around the diagonal. The protein prediction also shows a cluster of points (residues in positions 40-70 against residues around positions 100) that do not reflect any structural contact.

We tried to improve the prediction power of the analysis by increasing the number of mutations with successive generations of molecular evolution, but we observed only a strong enrichment of proximal interactions (near the diagonal of the contact map), at the expense of long range contacts. The strongest predictions from these mutational data appear to mimic the interactions observed during the early events of protein folding (Rose 1979), where the first and stronger connections are established between adjacent secondary structure elements. In general, the predicted contact distribution was non-random and contacts tended to crowd near the extremities of helices ignoring highly conserved areas like H2 (residues 67-83). Other minor crowding was observed around two big loop regions (90-100 and 160-170). The N-terminal crowding of contacts is likely the consequence of the degeneration and duplication around the starting site that was already observed during Sanger sequencing, while the C-terminal density is probably caused by sequential mutated positions in sequences where a C-terminal frameshift creates a block of strongly correlated positions without significantly affecting the functionality of the protein. The sparse number of contacts in conserved areas like H2 reflects the difficulty of creating a robust prediction when observing an inadequate number of mutations (**Figure 3A**). We thus face an interesting problem: the more contacts a residue is involved in, the more harmful a mutation becomes and we observe a limited number of variations. Since the mutational space at each position dictates the prediction power, the contacts formed by the most important residues will also be the ones harder to predict.

### Improving the prediction power in key areas and retrieval of long range interactions

To improve the accuracy and the spread of the predictions we applied a correlation-based approach identical to that proposed for fitness (Schmiedel and Lehner 2019). Residues in structural proximity are often deeply interconnected and likely to share the same environment.

Consequently they produce similar interaction patterns. Exploiting this similarity, we could obtain interactions from conserved positions by calculating partial correlation of the protein positions on the DCA patterns. This is because highly interconnected positions will have a characteristic association pattern across the protein easily recognizable by partial correlation, even if the original DCA predictions are inaccurate.

To validate this approach, we calculated the partial correlation with the UniProt dataset (**Suppl Figure S10**). As expected, the predictions from partial correlation are similar to the coupling scores obtained by DCA (**Figure 4C**) and are in general less densely packed around the diagonal albeit showing a few more incorrect predictions.

The partial correlation approach applied to the molecular evolution dataset gave very different results compared to the original coupling score (**Figures 4D**) and resulted more similar to what observed in the UniProt dataset, where the predicted interactions were more broadly distributed and both the terminals and the diagonal far less crowded (long range contacts (>50 residues): gen5: 55; gen12: 13; gen12 (partial correlation): 31; Uniprot: 71). The accuracy of the prediction was relatively low (**Suppl Figure S11**), even though several times bigger than the random expectation. This inaccuracy was caused by low precision and not by a low trueness to the underlying values as proven by the low value of shortest path from a true contact observed for the predicted pairs (**Suppl Figure S12**). Along the contact map diagonal we observed densities in correspondence to strong secondary structure interactions, like the proximity between N-terminal sheets and helix H1 represented in the graph by the cluster of contacts near residues 25 to 60. Other off-diagonal crowding (around residues 90-170) could be observed in the helical domain in correspondence to the interactions formed by the bending of the peptide chain in a turn. These interactions and similar ones, formed between the C-terminal half of the five stranded sheet and the C-terminal helices of the protein (200-285), were also visible in the original DCA score (**Figure 4B**) and the most evident areas along the diagonal of the UniProt dataset where the predicted interactions clustered (**Figure 4C**). Long range contacts, represented in the contact map by data points far from the diagonal, were significantly different if we evaluated the interactions obtained by partial correlation and those predicted by the original DCA score. Partial correlation prediction showed several off-diagonal prediction points, mainly associated with highly interconnected regions or between elements of the hydrophobic core. In particular, we observed several contacts of H2 (67-83) with other elements of the helical domain (H2 to H10, residues 65-210). This demonstrated the centrality of H2, even though the region is per se characterized by a small mutational landscape (**Figure 3A**). The analysis identified also another cluster in the helical domain describing the proximity of helix H10 (199-210) to helix H5 (117-126).

Overall, we were able to obtain a contact map that matches effectively that of the crystal structure without any prior structural information. Our analysis demonstrated the possibility to obtain evolutionary couplings from a collection of sequences evolved in vitro. DCA highlighted the strongest evolutionary signal of proximal interactions (around the diagonal of the contact map) while partial correlation extracted information on the relations between secondary structure elements. These results demonstrate that molecular evolution can be used as a powerful tool for structural prediction.

## Discussion

The study of evolutionary couplings is an emerging frontier of bioinformatics, able to retrieve the network of interactions that dictate protein fold and function (Weigt et al. 2009; Marks et al. 2011; Morcos et al. 2011; Ekeberg et al. 2013; Kamisetty et al. 2013; Ovchinnikov et al. 2014; Ovchinnikov et al. 2017). The innovation brought by the technique is the ability to produce structural information without the need of experimental structure determination, relying only on the traces left by evolution on protein sequence. The correlations are obtained from the continuous polishing process that the flow of time exerts on sequence to optimize/retain function. This makes any structural information retrieved by the analysis like a fossil imprint of an *in vivo* interaction.

The current computational techniques based on evolutionary couplings require thousands of sequences to provide statistically meaningful results (Morcos et al. 2011; Marks et al. 2012; Ekeberg et al. 2013). Thus, current evolutionary coupling methods are limited to ancient and universal protein families, for which sequence data are available across a huge variety of species. This is a major limitation: a large number of human proteins, for instance, do not have ancient phylogenetic origin (Lander et al. 2001). They are therefore not amenable to evolutionary coupling methods based on phylogenetic databases and can only be tackled by experimental approaches. Another advantage of mutational libraries with respect to the classical phylogenetic data is the representation of a sequence instead of a family, since the Markovian models that retrieve the sequences for the alignments in the standard analysis do not differentiate close paralogs from true orthologs. This poses a serious limitation for protein families rich in paralogs like globulins, for which it is nearly impossible to obtain information for a specific member of the family. The ability to represent a protein instead of a family is a new feature that can enable to distinguish a different level of details during the biological interpretation of the data.

Here, we presented a strategy (CAMELS) that overcomes this limitation and lays the bases to develop a general method to gather structural information on protein contacts without performing experimental structural studies or the need for thousands of natural variants of the target protein across natural evolution. We provided a unique pipeline from the molecular to the computational levels using most advanced techniques and solved a number of crucial technical problems. Because DCA is good at capturing compensating mutations, a high mutational load in the collection of functional sequence variants is recommended. When a single harmful mutation appears in the sequence, the protein will likely not be functional and bacteria that carry that specific variant will die. However, if a second mutation able to compensate the damage is also present in the sequence, the function of the protein can be restored and the host cell survives. At the time that the selection is introduced, both mutations must already be present in the sequence, hence the more mutations are inserted in each round of mutagenesis, the better. We favored this coincidence by increasing the selective pressure in the generations that we sequenced. This way, if a single mutation is harmful but still barely allows survival in a low selective pressure, there will still be few generations in which the second compensating mutation could occur before the strong selection of the last generation reaps all the sequences carrying mutations that are not compensated.

The CAMELS method is based on the power of phenotypic selection. We produced one of the largest and most diversified molecular evolution libraries that shows high single molecule sequencing quality and sequence divergence of nearly 10% (i.e. 25 amino acid mutations and around 55 mismatching nucleobases) from the original ancestral protein. It is also the first library to have been sequenced at the full-length protein level by third NGS. Other databases of TEM-1 mutagenic variants are available, some of which were created using epPCR (Jacquier et al. 2013) or deep mutational scanning (Firnberg et al. 2014) as the mutagenic mechanism. These public datasets cannot be used to infer structural information because they are mostly composed of single amino acid variants and thus cannot generate evolutionary coupling. The sequencing reads do also not always cover the full-length molecule thus losing long range information (Firnberg et al. 2014). Our method is different since it produces a deep, high quality and full-length sequencing of a prolonged selection-driven evolution of the TEM-1 lactamase instead of focusing on the effects of single mutations.

We used our library to obtain structural information, by creating sequence diversity through mutation and analysis of artificial evolutionary couplings. We showed that the predicted contact map matches successfully that of the reference crystal structure even though at the cost of a bias towards short and medium range contacts. These results show that we are effectively simulating the course of evolution even if we cannot entirely compress the millions of years of natural selection into the couple of months of in vitro mutagenesis and selection. We are nevertheless successfully following the early stages of the landscape exploration of the evolving protein using this to extract direct information about protein folding.

Pilot work on structure prediction from molecular evolution experiments have been published during the development of the present study (Figliuzzi et al. 2016; Rollins et al. 2019; Schmiedel and Lehner 2019). Our approach offers several advantages as compared to these methods. The strategies previously used can only be applied to proteins strictly under 200 amino acids and can thus be used solely on a small fraction of the proteome from all three domains of life (Zhang 2000). In particular, crucial for the success of our method is the growth of the library in a matrix of a semisolid medium, which allows local growth but prevents diffusion. Importantly, the use of third generation sequencing is a strong advantage of our method that can be used to easily overcome the sequence read length limitations of traditional sequencing platforms. A key advantage of CAMELS is the absence of protein length constraints, since both the mutagenesis strategy and the sequencing allow processing of proteins of any length.

Another limit of previous techniques is the impossibility to exhaustively sampling all double mutants in the limited libraries that can be screened in complex systems like human tissue cultures. Structure determination with the previous methods would only be achievable if the libraries were biased to massively reduce diversity. Therefore previous strategies are best suited to small proteins, or to protein systems where the directed evolution strategy can handle large libraries, such as in the case of the GB1 domain (Olson et al. 2014). CAMELS employs instead multiple rounds of mutation enrichment to compress the variability in a library of few hundred thousand elements. This solves the problem of limited library diversity and has the potential to be of practical value to investigations of moderately sized proteins in systems where the library size is an important constraint.

The CAMELS method is in principle generalizable: by generating hundreds of thousands of mutagenic functional variants, it permits to focus on any protein and builds the foundation for a targeted structural analysis. This may allow to investigate by DCA-like methods evolutionary younger proteins, like eukaryotic-only or vertebrate-only proteins or human proteins of neurobiological interest, ultimately solving species-specific questions that need species-specific answers.

What are the current limits of CAMELS? The most important one is the inability to fully reconstruct the protein structure in the current proof of concept formulation of the technology. Nevertheless CAMELS provides local and long range contacts. Improvements might be envisaged to overcome this limitation in future work. The main factors that could allow a complete structure determination are likely the number and distribution of mutations, the sequencing depth and the strength of the selective pressure. All these parameters can be easily scaled up, under straightforward conditions, that were, however, beyond the scope of the present proof-of-concept study.

The data revealed a strong correlation between the mean crystallographic B factor of residues in the reference structure and the information entropy retrieved from the evolved library. This happens because residues that are part of ordered regions as in the protein core are adverse to mutation and thus the most conserved ones. Harming these zones would affect the fold and function. A drastic increase in the number of mutations would help to generate variability in these conserved key residues that could be translated in better evolutionary couplings. The same logic applies by forcing a distortion in the mutation propensities to favor the generation of mutations in key areas. The problem of this approach is that we would distort the landscape obtained by unbiased epPCR. Modifying the driving force of the mutagenesis from epPCR to deep mutational scanning could provide a diverse landscape that might produce more precise couplings for the reasons mentioned. A critical comparison between the landscapes and couplings produced by epPCR and deep mutational scanning may be important to improve the technique in future implementations. Increasing the sequencing depth and the corresponding scale of selection is also an easy albeit currently expensive solution. This would likely increase the statistical power and allow low-entropy regions to show enough variation to be translated in couplings.

We noted that the fifth generation produced more long range contacts in the standard DCA prediction in respect to the twelfth. A possible explanation could be that the fifth generation produced nearly twice the number of reads of the twelve generation that in turn could alter the number of mutations in key areas. Another interpretation is that accumulation of mutations over the generations could create a broad compensatory effect that partially hinders the results. The correlated mutations accumulating throughout evolution are a mixture consisting of directly and indirectly interacting mutations. Since most random mutations are destabilizing, under the severe mutational load exerted on TEM-1 by selective pressure, most of the variants in the library should have a compromised stability. This strong selection pressure towards fixation of stabilizing mutations might thus favor the accumulation of correlated mutations of residues that do not directly interact. A more systematic investigation of this point using different target proteins will be important in the future.

An important requirement of CAMELS is phenotypic selection, a necessity that makes the method truly evolutionary: like in the natural environment, selection is always based on the target protein function. As a consequence, selection must be designed on a case-by-case basis. For some proteins (such as TEM1 beta lactamase), a phenotypic selection scheme is readily designed. More in general, selection schemes based on interactions could be considered that is probably the best approach to generalize the method. CAMELS could easily be modified, for instance, for the study of protein-protein interactions, exploiting selection schemes for interacting proteins coupled to SMRT sequencing, which would allow observation of protein pairs in a single sequencing read. Selection schemes based on signaling by the mutated target protein could also be envisaged.

The next obvious step will be to exploit standard and generic selection methods that rely on the folding and binding properties of the mutant proteins in the library, regardless of their functional activity. We have, for instance, already planned to use selection schemes to select for interacting partners, using a strategy we already pioneered for screening more stable antibodies against a given target (Visintin et al. 1999; Chirichella et al. 2017). We could apply CAMELS to two covariant interacting proteins, which could then be co-selected by a two-hybrid scheme for preserving their mutual binding. This strategy will provide information on the direct or indirect structural determinants for protein-protein interacting domains. This would be a revolutionary breakthrough that is not restricted to specific cases. It should be noted however that to successfully apply CAMELS, a good selection is critical. Although testing the function of a protein may seem a simple requirement on paper, the difficulty of developing an effective screening strategy cannot be underestimated especially for understudied proteins.

An elegant recent study successfully used deep mutagenesis to attempt determination of an unknown structure of a large complex human receptor in a physiologically relevant active conformation (Park et al. 2019). However, this example was somewhat limited to the specific structure of the protein. It will be interesting to apply CAMELS to members of the GPCR family, exploiting the signal transduction propriety of the receptors coupled to a screenable selection readout.

Finally, one of the biggest obstacles to an *in vitro* evolution approach was the precarious equilibrium between mutagenic strength and selection survival rate. We solved this issue with a generational approach. We can further envisage future applications of the method to a continuous evolution in a specialized bioreactor. Overall, the CAMELS method provides a solid methodology that bypasses the most limiting factors of evolutionary coupling analysis techniques and opens a new page in structural biology and evolution.

## Materials and methods

### Plasmid construction & cloning

The backbone plasmid vector pUC19 (Norrander et al. 1983) (ATCC 37254) from ThermoFisher Scientific (SD0061) was modified to add flanking XhoI and NheI restriction sites to the already present Amp^R^ ORF to be able to easily clone in later steps the mutagenized Amp^R^,. To construct the plasmid, both the β-lactamase gene and the complementary plasmid vector fragments were amplified with oligonucleotides carrying the XhoI and NheI restriction sites (XhoI_bla_fw: tgaaaactcgaggaagagtATGAGTATTCA, NheI_bla_rv: acttgggctagctctgacagTTACCAATGC; NheI_backbone_fw: gtcagagctagcccaagtttactcatatat, XhoI_backbone_rv: ctcttcctcgagttttcaatattattgaag). They were then digested with the restriction enzymes and ligated with T4 ligase (**Suppl Figure S12**). The 5’ restriction site was placed just behind the Shine-Dalgarno sequence and the ability to metabolize ampicillin was assessed by growth of *E.coli* carrying the plasmid in selective media. The new plasmid is named pUC19a. (**Suppl Figure S13**). The Amp^R^ gene of pUC19 (GenBank: M77789.2) expresses a TEM-1 (class A) β-lactamase whose structure can be viewed in the 1ZG4 PDB entry.

### Error prone PCR

Mutagenesis of the Amp^R^ gene was achieved with epPCR (Wilson and Keefe 2001) in a mutation prone buffer with manganese ions, low magnesium, unbalanced dNTPs concentrations and a low fidelity DNA polymerase. Both low magnesium and the presence of manganese ions affect the efficiency of magnesium ions as cofactors of the polymerase by competition or by sheer low availability, while the unbalanced dNTP concentration favors mutations by scarcity of substrate and the deliberate usage of a low fidelity polymerase further increases the mutation rate. The reaction mix contained Tris-HCl pH 8.3 10 mM, KCl 50 mM, MgCl2 7 mM, dCTP 1 mM, dTTP 1 mM, dATP 0.2 mM, dGTP 0.2 mM, 5’ primer (bla_mut_fw: tgaaaactcgaggaagagtATG) 2 μM, 3’ primer (bla_mut_rv: acttgggctagctctgacagTTA) 2 μM, template DNA 20 pg/μl, MnCl2 0.5 mM (added just before reaction starts), Taq G2 DNA polymerase (Promega M784A) 0.05 U/μl (added just before reaction starts). The error prone PCR was carried out in serial reactions of 4 cycles in 100 μL in the recommended supplier reaction conditions and with an annealing temperature of 62°C. In the first reaction tube, the DNA template was a gel purified XhoI/NheI digested β-lactamase fragment 20 pg/μl, while subsequent reactions were fed with 10 μL of the previous PCR product.

### Library construction

The purification and digestion protocols before library construction changed slightly among generations. However, the optimized version of the pipeline employed in the last generations proceeded as follows: gel purify ~80 μL of the PCR reaction mixture underwent a cumulative amount of 20 cycles of error prone PCR, avoiding carrying over other reaction byproducts as much as possible. PCR was performed in standard reaction conditions to amplify the product and guarantee that the two strands of the amplicons did not contain mismatching base pairs. This step helped reducing the ambiguity in base calling during the circular consensus analysis. The purified PCR product was digested with XhoI and NheI restriction enzymes for 3h in CutSmart buffer (NEB). One hour before the end of the reaction, an appropriate amount of calf intestinal phosphatase (CIP) (NEB M0290S) was added following the supplier’s instruction. Adding CIP during insert digestion strongly reduced the formation of insert concatemers, guaranteeing a single β-lactamase variant per plasmid. After gel purification to remove the CIP, the ligation between the fragment and the XhoI/NheI digested backbone of pUC19a was performed in a 1:1 insert:vector ratio. Formation of backbone concatemers was expected and unavoidable, but did not hinder the selection efficiency.

### Selection

The ligated library was purified and then transformed by electroporation in ElectroMAX DH5α-E competent cells (Invitrogen #11319019). We employed ultralow gelling agarose (SeaPrep, Lonza #50302) 0.3% in Luria Broth (LB) medium with ampicillin 25 μg/mL to grow the bacteria (Elsaesser and Paysan 2004; Fantini, Pandolfini, et al. 2017) obtaining between 0.4 and 3 million surviving colonies per liter. The bacterial growth in the fifth and twelfth generation was performed in LB medium with ampicillin 100 μg/mL to increase the stringency of the selection before the sequencing. After 40 h growth, the bacterial pellet was retrieved by centrifugation 7500RPM at RT and the plasmids extracted by a maxi prep.

### Sequencing

Construction of the libraries and sequencing on PacBio Sequel platform were carried out by Arizona Genomics Institute (AGI). After sequencing, the library was processed with the PacBio official analysis software SMRTlink to obtain the circular consensus (using ccs2) of the reads. In this step, the sequences where the consensus was built from less than 10 sequencing polymerase passes or when the predicted accuracy was less than 100 ppm (Phred 40) were filtered out from the dataset. The result was then mapped to the wild type β-lactamase XhoI-NheI digestion fragment of pUC19a with bowtie2 (Langmead and Salzberg 2012) to retrieve the coding strand and the start site of the lactamase. After *in silico* translating the dataset, protein collection was further refined keeping only the elements coding a protein of 286 amino acids (as the wild type) and then aligned using MAFFT (http://mafft.cbrc.jp/alignment/software/) (Katoh 2002) to construct the MSA. The 12th generation had issues with the *in silico* translation step caused by degeneration of the N-terminus as well as the starting site, resulting in a big amount of sequence with a premature termination codon. To circumvent the problem, the longest open reading frame, identified with a custom script, was considered the correct genetic sequence and the translated products were filtered to keep the sequences coding for proteins of at least the wild type length. This procedure was required to remove from the alignment bad quality reads, unrelated sequences and protein variants carrying a frameshift which would generate a strong correlation noise between adjacent amino acid positions. It is interesting to notice that classical evolutionary data supplied to the algorithm do not have this problem, and thus this is a new issue brought by the mutagenic data. This is probably because frame shifted sequences do not carry a sufficient neutrality to the system to be retained throughout natural evolution, while in the small landscape generated by the mutagenesis, every protein that satisfies the selection criteria of activity will be part of the collection.

To compare our data to the natural occurring mutations of TEM beta lactamase, we created a reference dataset by running a small seed of TEM beta lactamases in Hmmer (Finn et al. 2011) on the UniProt database (https://www.uniprot.org). An alternative dataset that could be used as a control is the Pfam family of beta-lactamase2 (PF13354). Since the Hmmer dataset from Uniprot is a collection of sequences that specifically matched the profile of the TEM family while Pfam family is a more general beta-lactamase collection we preferred the former as reference for our analysis.

### Direct Coupling Analysis

The predicted contact pairs were obtained using a custom implementation (Fantini, Malinverni, et al. 2017) of the asymmetric version of the DCA (Weigt et al. 2009; Morcos et al. 2011) that applies the Pseudo-likelihood method to infer the parameters of the Potts model (Balakrishnan et al. 2011; Ekeberg et al. 2013):

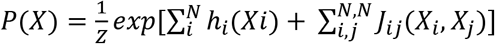

where X is a sequence of the MSA and Z is the partition function.

Sequences were reweighed using an identity threshold that reflects the mutation rate of the generation analyzed to remove parental inheritance (intended as “phylogenetic” bias created during mutagenesis) and sampling biases in the MSA. The first generation was too similar to the wild type to apply any sampling correction without unreasonably reducing the number of effectively non-redundant sequences (Morcos et al. 2011). The fifth generation used a 95% identity threshold and the twelfth generation a 90%. A standard L2 regularization was added following the original regularization described in Ekeberg et al., 2013 (Ekeberg et al. 2013) (λ = 0.01). The code used the scoring scheme for contacts proposed in Markley et al. (Markley et al., 2013) where the DCA scores were computed as the Frobenius norm of the local coupling matrices of the Potts model. Dunn et al. average product correction (APC) was subtracted to remove background correlation (Dunn et al. 2008). The N top scoring contact predictions (N equals the MSA sequence length) were compared with the contact map of the reference structure (1ZG4) constructed considering two residues to be in contact if at least a pair of their respective heavy-atom (non hydrogens) was less than 8.5 Å apart (Ekeberg et al. 2013). As it is standard practice, we removed predictions along the diagonal of the contact map if the residue pairs were less than five positions apart to promote enrichment of long-range predictions. Increasing this threshold to 8 or more did not change significantly the contact map prediction (data not shown). We used the shortest-path (SP) distance (Malinverni et al. 2015) defined as the L1 norm in the contact map lattice to join DCA predictions and the closest structural contact to visualize the agreement between predictions and empirical observations.

### Partial Correlation

Partial correlation is a measure of the correlations between two variables after removing from both the possible correlations they might have with another set of confounding variables. In other words partial correlation is the correlation between the residuals obtained from the regression of the variables of interest to the confounding variable set. In the present work, partial correlation is used to infer the correlation between each set of two rows of the DCA score matrix, removing the correlation these rows might have to all other rows of the matrix. Each row of the DCA matrix is a vector of the strength of association between the residue represented by the row and every other residues of the protein. Thus the partial correlation of the DCA matrix represents the similarity of the profile of these vectors, considering and removing the correlation with the profile of all the other residues. To obtain the partial correlation matrix from the symmetrical DCA score matrix, we first set all the diagonal elements of the matrix to 1 and then approximated the partial correlation between rows with the pcor.shrink function of the corpcor R package. The package implements a James-Stein estimator for the covariance matrix. The details of the method are explained in (Schäfer and Strimmer 2005) and in the manual of the package (https://cran.r-project.org/web/packages/corpcor/corpcor.pdf).

### Other bioinformatic tools

Graph generation was performed with R version 3.2.3 (2015-12-10). Poisson regression was performed with the fitdistrplus R package, while PCA was performed with the base R prcomp function. Mutation rates and Shannon information entropies were calculated with custom scripts. t-SNE was performed in Matlab version R2018b using the Hamming distance as metric.

## Acknowledgements

We thank our colleagues from Scuola Normale Superiore Federico Cremisi and Alessandro Cellerino for valuable comments. We are also grateful to Martina Goracci and Ottavia Vitaloni for times when, thanks to their wisdom and expertise, we were able to make major breakthroughs. We are also indebted to Duccio Malinverni for providing the software implementation of the asymmetric version of the DCA and to Alessandro Viegi for administrative support throughout the project. The project was funded by institutional funds from Scuola Normale Superiore, a BlueSky grant from University of Pavia and the UK Dementia Research Institute (RE1 3556) which is funded by the Medical Research Council, Alzheimer’s Society and Alzheimer’s Research UK.

## Data availability

All custom scripts used in this study can be accessed upon request. All sequencing data that support the findings of this study have been deposited in the National Centre for Biotechnology Information Sequence Read Archive (SRA) and are freely accessible through the SRA accession code PRJNA528665 (http://www.ncbi.nlm.nih.gov/sra/PRJNA528665).

The datasets of TEM-1 sequence variants collected in this study and the top contacts predicted in the analyses can be downloaded from the BioSNS website http://laboratoriobiologia.sns.it/supplementary-mbe-2019/

## Bibliography

Abraham EP, Chain E. 1940. An enzyme from bacteria able to destroy penicillin [1]. Nature 146:837.

Altschuh D, Lesk AM, Bloomer AC, Klug A. 1987. Correlation of co-ordinated amino acid substitutions with function in viruses related to tobacco mosaic virus. J. Mol. Biol. 193:693–707.

Ambler RP, Coulson AF, Frère JM, Ghuysen JM, Joris B, Forsman M, Levesque RC, Tiraby G, Waley SG. 1991. A standard numbering scheme for the class A beta-lactamases. Biochem. J. 276:269–270.

Balakrishnan S, Kamisetty H, Carbonell JG, Lee S-I, Langmead CJ. 2011. Learning generative models for protein fold families. Proteins 79:1061–1078.

Bershtein S, Segal M, Bekerman R, Tokuriki N, Tawfik DS. 2006. Robustness–epistasis link shapes the fitness landscape of a randomly drifting protein. Nature 444:929–932.

Bush K. 1997. Nomenclature of TEM beta-lactamases. J. Antimicrob. Chemother. 39:1–3.

Chen K, Arnold FH. 1993. Tuning the activity of an enzyme for unusual environments: sequential random mutagenesis of subtilisin E for catalysis in dimethylformamide. Proc. Natl. Acad. Sci. 90:5618–5622.

Chirichella M, Lisi S, Fantini M, Goracci M, Calvello M, Brandi R, Arisi I, D’Onofrio M, Di Primio C, Cattaneo A. 2017. Post-translational selective intracellular silencing of acetylated proteins with de novo selected intrabodies. Nat. Methods 14.

Deng Z, Huang W, Bakkalbasi E, Brown NG, Adamski CJ, Rice K, Muzny D, Gibbs RA, Palzkill T. 2012. Deep sequencing of systematic combinatorial libraries reveals β-lactamase sequence constraints at high resolution. J. Mol. Biol. 424:150–167.

van Dijk EL, Jaszczyszyn Y, Naquin D, Thermes C. 2018. The Third Revolution in Sequencing Technology. Trends Genet. 34:666–681.

Dunn SD, Wahl LM, Gloor GB. 2008. Mutual information without the influence of phylogeny or entropy dramatically improves residue contact prediction. Bioinformatics 24:333–340.

Eid J, Fehr A, Gray J, Luong K, Lyle J, Otto G, Peluso P, Rank D, Baybayan P, Bettman B, et al. 2009. Real-Time DNA Sequencing from Single Polymerase Molecules. Science 323:133–138.

Ekeberg M, Lövkvist C, Lan Y, Weigt M, Aurell E. 2013. Improved contact prediction in proteins: Using pseudolikelihoods to infer Potts models. Phys. Rev. E - Stat. Nonlinear, Soft Matter Phys. 87.

Elsaesser R, Paysan J. 2004. Liquid gel amplification of complex plasmid libraries. Biotechniques 37:200–202.

Fantini M, Malinverni D, De Los Rios P, Pastore A. 2017. New techniques for ancient proteins: Direct coupling analysis applied on proteins involved in iron sulfur cluster biogenesis. Front. Mol. Biosci. 4.

Fantini M, Pandolfini L, Lisi S, Chirichella M, Arisi I, Terrigno M, Goracci M, Cremisi F, Cattaneo A. 2017. Assessment of antibody library diversity through next generation sequencing and technical error compensation. Budak H, editor. PLoS One 12:e0177574.

Figliuzzi M, Jacquier H, Schug A, Tenaillon O, Weigt M. 2016. Coevolutionary landscape inference and the context-dependence of mutations in beta-lactamase tem-1. Mol. Biol. Evol. 33.

Finn RD, Clements J, Eddy SR. 2011. HMMER web server: Interactive sequence similarity searching. Nucleic Acids Res. 39.

Firnberg E, Labonte JW, Gray JJ, Ostermeier M. 2014. A Comprehensive, High-Resolution Map of a Gene’s Fitness Landscape. Mol. Biol. Evol. 31:1581–1592.

Göbel U, Sander C, Schneider R, Valencia A. 1994. Correlated mutations and residue contacts in proteins. Proteins Struct. Funct. Genet. 18:309–317.

Goldsmith M, Tawfik DS. 2009. Potential role of phenotypic mutations in the evolution of protein expression and stability. Proc. Natl. Acad. Sci. 106:6197 LP–6202.

Hopf TA, Colwell LJ, Sheridan R, Rost B, Sander C, Marks DS. 2012. Theory Three-Dimensional Structures of Membrane Proteins from Genomic Sequencing. Cell 149:1607–1621.

Hopf TA, Schärfe CPI, Rodrigues JPGLM, Green AG, Kohlbacher O, Sander C, Bonvin AMJJ, Marks DS. 2014. Sequence co-evolution gives 3D contacts and structures of protein complexes. Elife 3:e03430.

Huang W, Palzkill T. 1997. A natural polymorphism in β-lactamase is a global suppressor. Proc. Natl. Acad. Sci. 94:8801 LP–8806.

Jacquier H, Birgy A, Le Nagard H, Mechulam Y, Schmitt E, Glodt J, Bercot B, Petit E, Poulain J, Barnaud G, et al. 2013. Capturing the mutational landscape of the beta-lactamase TEM-1. Proc. Natl. Acad. Sci.

Jelsch C, Mourey L, Masson J-M, Samama J-P. 1993. Crystal structure of Escherichia coli TEM1 β-lactamase at 1.8 Å resolution. Proteins Struct. Funct. Bioinforma. 16:364–383.

Kamisetty H, Ovchinnikov S, Baker D. 2013. Assessing the utility of coevolution-based residue-residue contact predictions in a sequence- and structure-rich era. Proc. Natl. Acad. Sci. U. S. A. 110:15674–15679.

Katoh K. 2002. MAFFT: a novel method for rapid multiple sequence alignment based on fast Fourier transform. Nucleic Acids Res. 30:3059–3066.

Kircher M, Stenzel U, Kelso J. 2009. Improved base calling for the Illumina Genome Analyzer using machine learning strategies. Genome Biol. 10:R83.

Lander ES, Linton LM, Birren B, Nusbaum C, Zody MC, Baldwin J, Devon K, Dewar K, Doyle M, Fitzhugh W, et al. 2001. Initial sequencing and analysis of the human genome. Nature 409:860–921.

Langmead B, Salzberg SL. 2012. Fast gapped-read alignment with Bowtie 2. Nat. Methods 9:357–359.

Malinverni D, Marsili S, Barducci A, de Los Rios P. 2015. Large-Scale Conformational Transitions and Dimerization Are Encoded in the Amino-Acid Sequences of Hsp70 Chaperones. PLoS Comput. Biol. 11:1–15.

Marks DS, Colwell LJ, Sheridan R, Hopf TA, Pagnani A, Sander C. 2011. Protein 3D Structure Computed from Evolutionary Sequence Variation. 6.

Marks DS, Hopf T a, Sander C. 2012. Protein structure prediction from sequence variation. Nat. Biotechnol. 30:1072–1080.

Morcos F, Pagnani A, Lunt B, Bertolino A, Marks DS, Sander C, Zecchina R, Onuchic JN, Hwa T, Weigt M. 2011. Direct-coupling analysis of residue coevolution captures native contacts across many protein families. Proc. Natl. Acad. Sci. U. S. A. 108:E1293–301.

Norrander J, Kempe T, Messing J. 1983. Construction of improved M13 vectors using oligodeoxynucleotide-directed mutagenesis. Gene 26:101–106.

Olson CA, Wu NC, Sun R. 2014. A Comprehensive Biophysical Description of Pairwise Epistasis throughout an Entire Protein Domain. Curr. Biol. 24:2643–2651.

Ovchinnikov S, Kamisetty H, Baker D. 2014. Robust and accurate prediction of residue-residue interactions across protein interfaces using evolutionary information. Elife 2014:1–21.

Ovchinnikov S, Park H, Varghese N, Huang P-S, Pavlopoulos GA, Kim DE, Kamisetty H, Kyrpides NC, Baker D. 2017. Protein structure determination using metagenome sequence data. Science 355:294 LP–298.

Park J, Selvam B, Sanematsu K, Shigemura N, Shukla D, Procko E. 2019. Structural architecture of a dimeric class C GPCR based on co-trafficking of sweet taste receptor subunits. J. Biol. Chem. 294:4759–4774.

Pazos F, Helmer-Citterich M, Ausiello G, Valencia a. 1997. Correlated mutations contain information about protein-protein interaction. J. Mol. Biol. 271:511–523.

Rollins NJ, Brock KP, Poelwijk FJ, Stiffler MA, Gauthier NP, Sander C, Marks DS. 2019. Inferring protein 3D structure from deep mutation scans. Nat. Genet. 51:1170–1176.

Rose GD. 1979. Hierarchic organization of domains in globular proteins. J. Mol. Biol. 134:447–470.

Salverda MLM, De Visser JAGM, Barlow M. 2010. Natural evolution of TEM-1 β-lactamase: experimental reconstruction and clinical relevance. FEMS Microbiol. Rev. 34:1015–1036.

Schäfer J, Strimmer K. 2005. A Shrinkage Approach to Large-Scale Covariance Matrix Estimation and Implications for Functional Genomics. Stat. Appl. Genet. Mol. Biol. 4.

Schlessinger A, Rost B. 2005. Protein flexibility and rigidity predicted from sequence. Proteins Struct. Funct. Bioinforma. 61:115–126.

Schmiedel JM, Lehner B. 2019. Determining protein structures using deep mutagenesis. Nat. Genet. 51.

Stec B, Holtz KM, Wojciechowski CL, Kantrowitz ER. 2005. Structure of the wild-type TEM-1 β- lactamase at 1.55 Å and the mutant enzyme Ser70Ala at 2.1 Å suggest the mode of noncovalent catalysis for the mutant enzyme. Acta Crystallogr. Sect. D Biol. Crystallogr. 61:1072–1079.

Stiffler MA, Hekstra DR, Ranganathan R. 2015. Evolvability as a Function of Purifying Selection in TEM-1 β-Lactamase. Cell 160:882–892.

Toth-Petroczy A, Palmedo P, Ingraham J, Hopf TA, Berger B, Sander C, Marks DS. 2016. Structured States of Disordered Proteins from Genomic Sequences. Cell 167:158–170.e12.

Uguzzoni G, John Lovis S, Oteri F, Schug A, Szurmant H, Weigt M. 2017. Large-scale identification of coevolution signals across homo-oligomeric protein interfaces by direct coupling analysis. Proc. Natl. Acad. Sci. U. S. A. 114:E2662–E2671.

Visintin M, Tse E, Axelson H, Rabbitts TH, Cattaneo A. 1999. Selection of antibodies for intracellular function using a two-hybrid in vivo system. Proc Natl Acad Sci U S A 96:11723–11728.

De Visser JAGM, Salverda MLM, Barlow M. 2010. Natural evolution of TEM-1 β-lactamase: experimental reconstruction and clinical relevance. FEMS Microbiol. Rev. 34:1015–1036.

Wang B, Kennedy MA. 2014. Principal components analysis of protein sequence clusters. J. Struct. Funct. Genomics 15:1–11.

Wang X, Minasov G, Shoichet BK. 2002. The Structural Bases of Antibiotic Resistance in the Clinically Derived Mutant β-Lactamases TEM-30, TEM-32, and TEM-34. J. Biol. Chem. 277:32149–32156.

Weigt M, White R a, Szurmant H, Hoch J a, Hwa T. 2009. Identification of direct residue contacts in protein-protein interaction by message passing. Proc. Natl. Acad. Sci. U. S. A. 106:67–72.

Wilson DS, Keefe AD. 2001. Random Mutagenesis by PCR. In: Current Protocols in Molecular Biology. Vol. 51. Hoboken, NJ, USA: John Wiley & Sons, Inc. p. 8.3.1–8.3.9.

Zaccolo M, Gherardi E. 1999. The effect of high-frequency random mutagenesis on in vitro protein evolution: a study on TEM-1 beta-lactamase. J Mol Biol 285:775–783.

Zhang J. 2000. Protein-length distributions for the three domains of life. Trends Genet. 16:107109.

